# Odad1 is Required for Deployment of Axonemal Outer Arm Dyneins and Embryonic Left-Right Patterning

**DOI:** 10.64898/2026.10.19.760162

**Authors:** Nayeli G. Reyes-Nava, Indrani Basu, Kristen R. Holladay, John B. Wallingford

## Abstract

Motile cilia generate fluid flow required for development and tissue homeostasis. Defects in ciliary motility cause Primary Ciliary Dyskinesia (PCD), presenting with situs inversus, infertility, chronic respiratory disease, and hydrocephalus. Ciliary beating requires outer dynein arms (ODAs), anchored to microtubules by the ODA docking complex (ODA-DC). ODAD1 is an ODA-DC component and a known PCD protein. ODAD1 alleles are often associated with human laterality defects, however Odad1 has not been reported to localize to 9+0 axonemes. Here, we show that GFP-Odad1 localizes to nodal cilia (9+0) in the gastrocoel roof plate (GRP), the frog’s equivalent to the mammalian node where left-right asymmetry is first established. We found that Odad1 knockdown (KD) disrupts left-right patterning in *Xenopus*, with loss of *pitx2* expression and abnormal gut looping reflecting defective 9+0 nodal cilia at the GRP. In 9+2 multiciliated cells (MCC) axonemes, Odad1 loss impaired deployment of ODA motors and ODA-DC subunits, disrupting ciliary motility. Two patient-derived truncating *ODAD1* alleles were similarly unstable, failing to rescue ODA docking in MCCs or left-right patterning at the GRP. Together, these results show ODAD1 is required for ODA-DC assembly in both 9+0 and 9+2 cilia, linking its loss to ciliary motility and left-right patterning defects in ODAD1-associated ciliopathy.

## Introduction

Motile cilia drive fluid flow across vertebrate tissues, from the mucociliary epithelium that clears the airway to the rotational beating that breaks symmetry at the left-right organizer (LRO). The structural composition of motile cilia differs by tissue: airway cilia have a 9+2 microtubule arrangement, whereas nodal cilia are 9+0, lacking the central pair of microtubules — an absence thought to underlie their rotational, rather than planar, beating pattern [1], [2]. Both ciliary types depend on axonemal dynein arms, ATPase-driven protein motors that decorate the outer doublet microtubules of the axoneme, however the specific dynein subunits employed differ between ciliary types [3], [4], [5], [6], [7]. Outer dynein arms (ODAs) regulate the ciliary beating, and loss of ODAs is the leading cause of primary ciliary dyskinesia (PCD), a genetic heterogenous disorder characterized by chronic sino-pulmonary disease, infertility and laterality defects [8], [9], [10].

Proper ODA attachment to microtubules requires a specialized protein complex, the outer dynein arm docking complex (ODA-DC), which anchors ODAs to the microtubule doublet at defined periodic sites. Foundational work in *Chlamydomonas* first defined this complex biochemically, showing that a heterotrimeric docking complex — including a ∼70 kDa subunit later named DC70/Oda1 — is necessary and sufficient to tether ODAs to the axoneme, and that a single truncating mutation in Oda1 subunit abolishes ODA binding entirely [11], [12]. More recently, cryo-electron tomography revealed that the mammalian ODA-DC is more elaborate than its algal counterpart, comprising five subunits (ODAD1–ODAD5) rather than three, with ODAD1 forming part of the core scaffold onto which ODAD2, ODAD3, ODAD4, and ODAD5 hierarchically assemble [13], [14], [15], [16], [17].

The human ortholog of *ODAD1* (formerly *CCDC114*), was identified a decade later as a PCD-causing gene through independent exome sequencing [18], [19]. Both reports showed that loss-of-function variants in *ODAD1* cause loss of ODAs, resulting in immotile respiratory cilia, with affected patients frequently presented with situs inversus or other laterality defects [19]. A growing allelic series — including hypomorphic and splice-altering variants associated with intermediate or unusually mild phenotypes — shows that the cellular and clinical consequences of *ODAD1* dysfunction lie on a spectrum rather than a simple null/wild-type binary [20], [21], [22], [23], [24], [25].

*ODAD1* function has been increasingly investigated in recent years, from initial reconstitution and genetics in a unicellular ciliated algae [11], [26], to genotype-phenotype correlation in human patient tissue [19], [21], [22], [23] and, more recently, patient derived airway organoids [20]. ODA-DC subunits ODAD2 (ARMC4), ODAD3 (CCDC151), ODAD4 (TTC25), and ODAD5 (CALAXIN) have been previously validated in animal models [14], [15], [27], [28]. Zebrafish knockdown of *armc4* causes dose-dependent heart-looping and left-right asymmetry defects [14]. Knockdown of *ccdc151* causes situs randomization, dilated pronephros, kidney cysts, and hydrocephalus, with immotile cilia and loss of ODAs in zebrafish [15]. Finally, knockdown of *ttc25* in *Xenopus* multiciliated cells disrupts ciliary beating and ODA assembly, while a CRISPR/Cas9 *Ttc25* knockout mouse independently shows laterality defects, immotile nodal cilia with absent leftward flow, and loss of ODAs by electron microscopy [27]. However, Odad1 function has not been assessed in *Xenopus*, or in any other animal model; and despite its implications in human laterality defects, no study has yet reported Odad1 to 9+0 axonemes. The mechanisms underlying ODAD1-associated PCD therefore remain incompletely understood.

Here, we investigated the mechanisms underlying Odad1 function, by exploring how loss of Odad1 disrupts ciliary assembly and function across distinct motile ciliated tissues. We show that of Odad1 localizes to nodal cilia axonemes, and that Odad1 loss disrupts ODA and ODA-DC subunits deployment to motile cilia, both nodal cilia and multiciliated cells (MCCs) cilia. These phenotypes later translate into developmental defects such as abnormal gut looping and deficient fluid flow. Additionally, we functionally tested uncharacterized patient-derived Odad1 alleles. The phenotypes resulting from Odad1 deficiency in *Xenopus* motile cilia, recapitulate human phenotypes associated with PCD and we confirm that tested variants correspond to loss-of-function alleles.

## Methods

### Animal husbandry

*Xenopus* experiments were conducted in accordance with the animal protocol AUP-2024-00130 and the animal ethics guidelines of the University of Texas at Austin. Ovulations were induced in female adult *Xenopus laevis* by injection of hCG (human chorionic gonadotropin).

### Microinjection of morpholino oligonucleotides and mRNA and embryo manipulations

*Xenopus laevis* embryos were obtained via *in vitro* fertilization using standard methods and staged according to Nieuwkoop and Faber [29]. Embryos were raised to the appropriate developmental stages in 1/3x Marc’s Modified Ringer’s (MMR).

Morpholino oligonucleotides and/or mRNAs were injected into four cell stage embryos. Ventral blastomeres were injected to target the multiciliated epidermis [30] and dorsal blastomeres at the dorsal marginal zone were injected to target nodal cilia in the *Xenopus* GRP as previously described [31].

A splice-site morpholino was designed to target the first exon-intron splicing junction of the of o*dad1* in the allotetraploid genome of *Xenopus laevis.* The splice blocking morpholino for *odad1* was injected at the concentration of 10 ng (5’ AATGCTGAAAATTACCTATTCCTTC 3’).

*Xenopus* Odad1 sequence was obtained from Xenbase (http://www.xenbase.org). Odad1 open reading frame (ORF) was amplified from the *Xenopus laevis* cDNA library by polymerase chain reaction (PCR) and then inserted into a pCS10R vector containing a fluorescent tag. In addition to the vectors described previously [30], [32], [33], the following constructs were cloned into pCS10R vector: GFP-Odad1, Myc-Odad1. To generate a *Xenopus* allele corresponding to the human patient allele, mutagenesis was performed on the GFP-Odad1, using Q5 Site-Directed Mutagenesis Kit (NEB, Cat #E0554S). All constructs were linearized and the capped mRNAs were synthesized using mMESSAGE mMACHINE SP6 transcription kit (ThermoFisher Scientific, Cat #AM1340). Each mRNA was injected into two ventral blastomeres at the four-cell stage (∼25-80 pg per blastomere).

Whole-mount *in situ* hybridization was performed as described [34]. Embryos were fixed in MEMFA for 1–2 hr and processed following standard protocols. Digoxigenin-labeled antisense RNA probes (Roche) were synthesized from linearized plasmid templates using SP6 RNA polymerase (Promega). Hybridization was performed against a *pitx2c* probe [35]. Stained embryos were imaged by brightfield microscopy at stage 28, and *pitx2c* expression was scored as normal, inverted, bilateral, or absent.

### RT-PCR

RT-PCR was performed to confirm the efficacy of Odad1 morpholino. Morpholino was injected into all blastomeres at four-cell stage of embryos. Toral RNA was isolated with Trizol reagent (Invitrogen, 15596026) from *N*=4 embryos at stage 26, and cDNA was synthesized with an M-MLV reverse transcription kit (Invitrogen, 28025013). *Odad1* and *odc1* were amplified by Taq-polymerase (Invitrogen, 10342020) with the following primers: *odad1 25* 5’-agtgcccgaagtgacagtag-3’, *odad1* 475 5’-cctgtcgagtctgttctcca-3’, and *odc1* primers were *odc1* 426F 5’-ggcaaggaatcacccgaatg-3’ and *odc1* 843R 5’-ggcaacatagtatctcccaggctc-3’.

### Gut looping

Embryos were injected at four-cell stage, targeting the GRP nodal cilia as above described. Gut morphology was scored at end stage (NF 45-46). The independent experiments from different clutches of embryos were performed. The data were then used to calculate the average percent of injected embryos with abnormal gut morphology from the three experiments (total control N=77; Odad1 KD N=65).

### Western blotting

Protein for western blotting was isolated from embryos (N=30 stage NF 25) were lysed in lysis buffer (ChromoTek, gtak-20) supplemented with protease inhibitors. The lysates were centrifuged to remove cell debris, and the supernatants were subjected to SDS-PAGE followed by immunoblotting using standard protocols. The antibodies used were as follows: 1:200 anti-GFP antibody (Santa Cruz Biotechnology, sc-9996), 1:2000 HRP-conjugated goat anti-mouse IgG (H+L) secondary antibody (Thermo Fisher Scientific, 31430), 1:10,000 anti-β-actin monoclonal antibody (Proteintech, 6009-1) and anti-Myc antibody (9E10; Abcam, ab32).

### Live imaging and image analysis

Live imaging of multi-ciliated cells was performed in *Xenopus* embryos expressing tagged proteins at stages 24–26. Live imaging of nodal cilia was performed from stage 18 dorsal explants as previously described [31]. Whole embryos or explants were mounted between coverslips and immersed in 1/3X MMR prior to imaging. Imaging was conducted using a Zeiss LSM700 laser scanning confocal microscope equipped with a Plan-Apochromat 63×/1.4 NA oil immersion objective (Zeiss) or a Nikon Eclipse Ti confocal microscope with a 63×/1.4 NA oil immersion objective. Quantitative image analysis was performed using Fiji. Graphs were generated, and statistical analyses, including error bars representing mean ± SD and P values, were conducted using Prism 10 software. Statistical significance was determined using a non-parametric Mann-Whitney U test for two-group comparisons and a one-way ANOVA for comparisons involving more than two groups.

### Time lapse movies (bead flow) and analysis

*Xenopus* embryos were grown up to stage 28 and mounted onto an imaging chamber in 1/3X MMR with green fluorescent beads (Fluospheres^TM^ polystyrene, 1.0 μm, yellow-green 505/515). Fluorescent bead flow was imaged in NF st. 28 *X. laevis* embryos using Nikon W1 Spinning Disc Confocal, acquiring 5 s continuous time-lapse videos (10 ms exposure, no interframe delay, 12-bit depth). Bead trajectories over 60 frames were visualized using Fiji’s Temporal-Color Code function (Image > Hyperstacks > Temporal-Color Code), mapping frame number to an Fire color scale from frame 0 (Blue) to frame 60 (white).

## Results

### Odad1 is important for establishing left-right asymmetry in *Xenopus*

Left-Right (LR) patterning in vertebrates is established by leftward fluid flow generated by motile cilia at the embryonic LRO, which triggers asymmetric expression of *nodal* pathway components, including *pitx2*, in the left lateral plate mesoderm [36], [37]. Given the role of Odad1 as an ODA-DC component required for ciliary motility and its association with LR defects in humans, we asked whether Odad1 is required for LR asymmetry in vertebrates.

LR patterning defects frequently manifest downstream organ laterality defects, so we examined gut looping in Odad1 deficient *Xenopus* embryos. An antisense morpholino oligonucleotide designed to block *Odad1* splicing severely depleted the mRNA as indicated by reverse transcription PCR (Fig. S1). Control embryos (NF st. 45-46) consistently displayed normal, stereotyped gut looping (Fig. 1A), whereas a significant number of Odad1 knockdown (KD) embryos exhibited disrupted gut looping (Fig. 1B,C). Interestingly, malrotations were not biased, but instead were randomized with the predominant phenotype being disrupted looping, suggesting a failure to establish the LR axis rather than a simple inversion of it.

**Figure 1.**
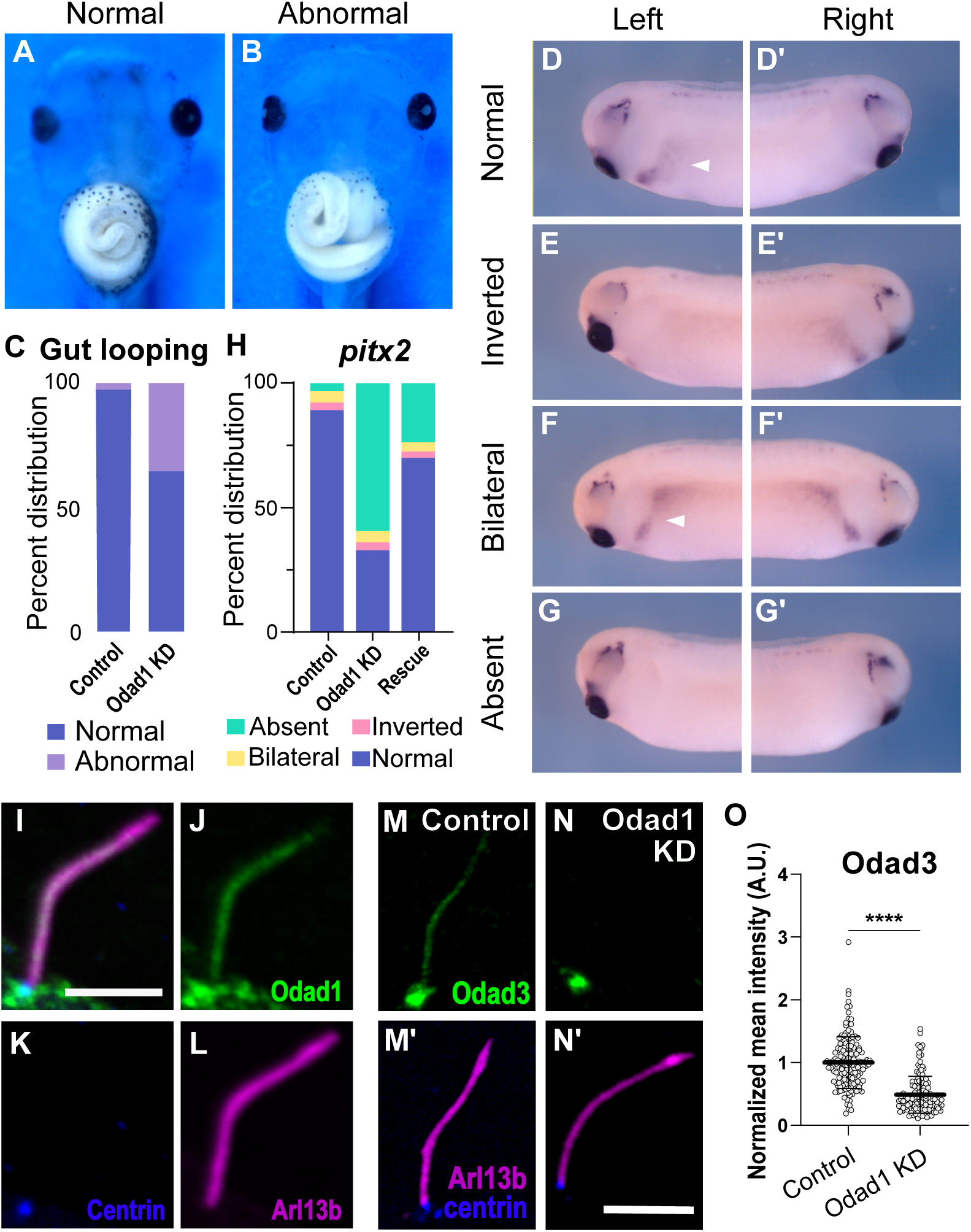
Loss of Odad1 results in left-right asymmetry defects. (A) Normal gut looping in a control stage 46 embryo (B) Gut looping is disrupted following Odad1 knockdown. (C) Percentage of normal or abnormal gut looping shown in control N=77 and Odad1 KD embryos N=65. (D-G) Whole mount in situ hybridization of left-right patterning marker *pitx2* in stage 28 *X. laevis* embryo imaged by brightfield microscopy. Colorimetric staining for *pitx2* RNA (purple) showing normal left side expression (D, D’), inverted (E,E’), bilateral (F,F’) or absent (G, G’) *pitx2* expression. White arrows pointing to *pitx2* expression in the lateral plate. (H) Percent distribution of *pitx2* expression in control N=63, Odad1 knockdown (KD) N=63, and rescue embryos N=80 across four independent experiments. (I-L) GFP-Odad1 localization pattern in nodal cilia, displaying axonemal localization as well as localization to the ciliary base. (I) Merged projection with Odad1 in green, Centrin-BFP labels basal bodies (K) and Arl13b is used as ciliary marker (L). (M-N) Odad1 knockdown results in decreased ciliary localization of Odad3 to nodal cilia. (O) Quantification of normalized Odad3 mean intensity along nodal cilia, showing decreased GFP signal in Odad1 KD embryos compared to controls.

Given that Odad1 loss disrupted the normal leftward looping of the gut, we next asked whether this reflected a broader failure to establish left-right asymmetry at the molecular level. We therefore examined expression of *pitx2,* the conserved downstream effector of left-right patterning, normally restricted to the left lateral plate mesoderm [38], [39], [40]. Using whole-mount in situ hybridization, we found that control embryos displayed the expected normal left-sided *pitx2* expression through stage 28 *Xenopus* embryos (Fig. 1D,D′). Interestingly, rather than shifting to the right side as would be expected for reversal, *pitx2* expression was lost altogether in most Odad1 KD embryos (>60% absent) (Fig. 1G,G’). Quantification across four independent experiments confirmed a shift in the distribution of *pitx2* expression patterns in Odad1 KD embryos (30% normal distribution), compared to controls (90% normal left expression) (Fig. 1 H). The absent *pitx2* phenotype was substantially rescued by co-injection of Myc-Odad1 mRNA (70% normal distribution) (Fig. 1G), confirming the specificity of the knockdown phenotype. Together, these results indicate that Odad1 is required for the establishment — not merely the orientation — of LR asymmetry in vertebrates.

We next sought to determine how Odad1 loss disrupts LR asymmetry at the cellular level, beginning with its subcellular localization in nodal cilia of the *Xenopus* gastrocoel roof plate (GRP), the amphibian equivalent of the mammalian LRO. GFP-Odad1 localized along the length of nodal cilia axonemes (Fig. 1I,J), labeled with the ciliary membrane marker Arl13b (Fig. 1L). In addition, GFP-Odad1 was enriched at the ciliary base, where it colocalized and extended around Centrin-BFP-labeled basal bodies (Fig. 1K). The subcellular localization of ODAD1 in the mammalian node remains unreported. Odad1 localization to nodal cilia axonemes is consistent with the canonical role in ODA docking across motile cilia, while its enrichment to the ciliary base is distinct from that reported in mammalian MCCs [13]. Surprisingly, previous reports using epitope-tagged Odad1 and immunofluorescence described Odad1 as restricted to the ciliary base of 9+0 cilia in the zebrafish KV and spinal canal [32,33].

Odad1 localization to nodal cilia led us to ask whether Odad1 loss affects the ciliary localization of other ODA-DC components. Similarly to Odad1, Odad3 localizes along nodal cilia axoneme and basal body (Fig. 1M,M’). We found that Odad1 KD resulted in a marked decrease in ciliary localization of GFP-Odad3 along nodal cilia (Fig. 1N,N’) compared to controls. Quantification of Arl13b normalized Odad3 mean fluorescence intensity along the ciliary axoneme confirmed a significant reduction in Odad3 signal in Odad1 KD embryos relative to controls (Fig. 1O). Together, these data indicate that Odad1 is required for proper docking or stabilization of Odad3 within nodal cilia, providing a mechanistic basis for the ciliary motility and LR patterning defects observed upon Odad1 loss.

### Odad1 is required for deployment of ODAs to multiciliated cells axonemes

In the LR organizer rotary beating of motile mono cilia (9+0) generates a leftward flow of extracellular fluid that establishes to the LR asymmetry of organs during development [41], [42]. In contrast, the power stroke from 9+2 cilia of multiciliated cells generates directional fluid flow, essential for mucociliary clearance in the airways [1]. This difference in beat pattern reflects their distinct axonemal architecture. Despite these structural and functional differences, both types of cilia depend on axonemal dyneins (ODAs) to generate the forces that drive cilia beating [1,3–5].

To better understand Odad1 functions, we investigated its role in *Xenopus* MCCs [43]. We first assessed fluid flow, the functional output of MCC cilia beating, in Odad1 KD embryos. Using spinning disk confocal microscopy, we imaged fluorescent tracer bead movement across the *Xenopus* embryo medium. Flow trace analysis revealed that in control embryos, bead trajectories aligned fluid flow near the embryo surface with less robust but still obvious flow in regions farther from the surface (Fig. S2A). Control embryo bead trajectories are consistent with a directional, robust, spatially coordinated ciliary-driven flow. By contrast, in Odad1 KD embryos, flow was markedly reduced, where trajectories were shorter and no clear flow in regions farther from the surface (Fig. S2B). These results are consistent with the reduced or abnormal ciliary beating previously reported in MCC cultures and organoids derived from ODAD1 patients [18], [19], [20], [23].

Because MCC beating is driven by ODAs, we investigated the effect of Odad1 KD in deployment of ODAs to MCCs axonemes. Odad1 knockdown disrupted the normal axonemal distribution of the ODA intermediate chain subunit Dnai2 (GFP-Dnai2) (Fig. 2B,B’), which is typically enriched along the length of the axoneme (Fig. 2A, A’). This defect was rescued by re-expression of Myc-tagged Odad1 in a dose dependent fashion (Fig. 2C-D, C’-D), demonstrating the specificity of the KD. Quantification—normalized to a membrane-RFP marker—confirmed a significant reduction in Dnai2 levels in knockdown MCCs that was restored upon rescue (Fig. 2E). To confirm this result, we examined an additional ODA subunit, the light chain ODA subunit Dnal4. We observed a similar phenotype, with Odad1 knockdown decreasing GFP-Dnal4 axonemal signal and rescue restoring it (Fig. 2F-H).

**Figure 2.**
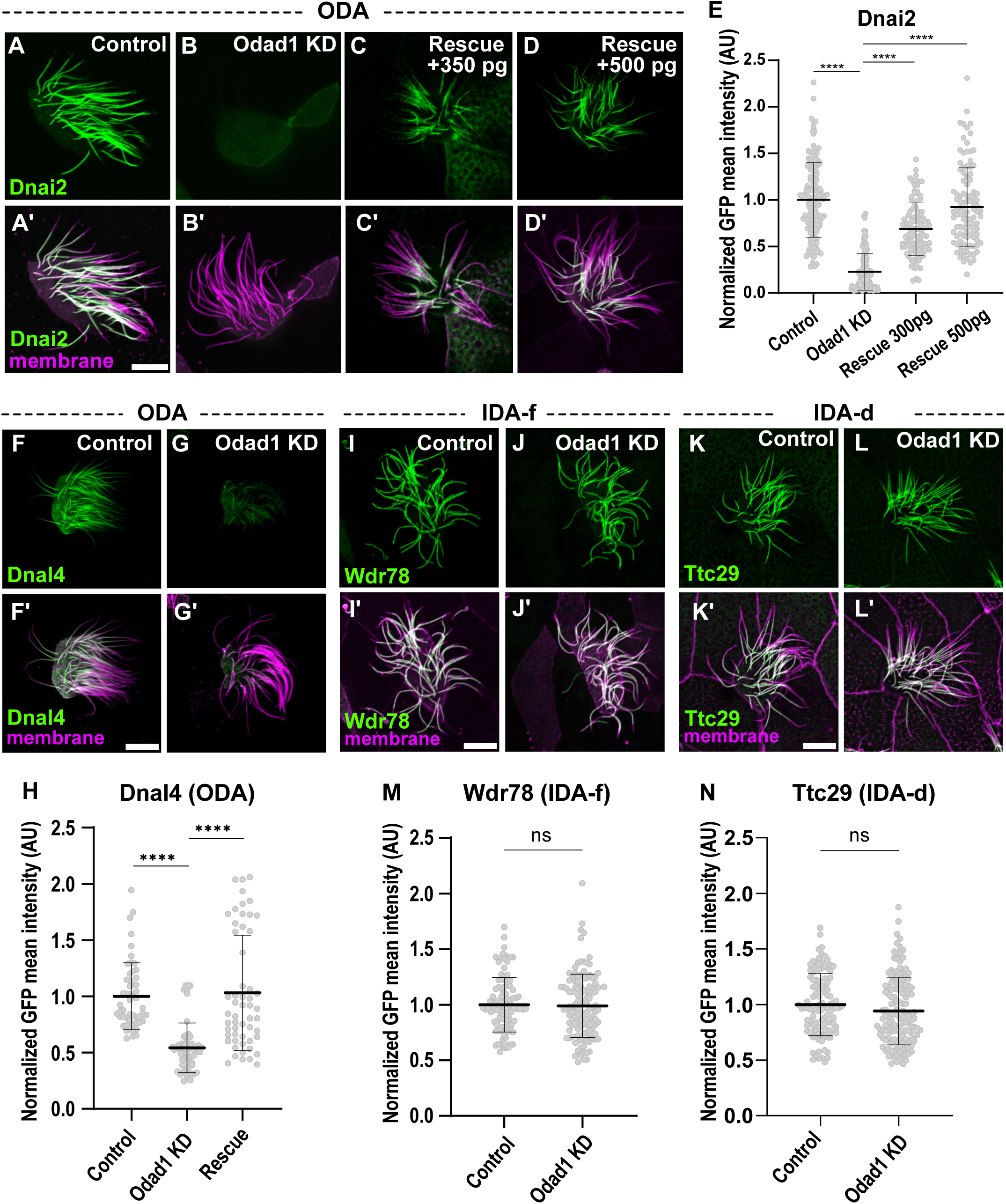
Odad1 knockdown results in loss outer dynein arms in MCCs. (A) GFP-Dnai2 labeling of motile axonemes in control MCCs shows normal localization ODA subunits. (A′) Merged channels (membrane labeled with CAAX-RFP, magenta and Dnai2, green) reveal normal cilia morphology in control MCCs. (B) GFP-Dnai2 is absent from motile cilia in Odad1 knockdown (KD) MCCs. (B′) Membrane labeling (magenta) in merged channels shows normal motile cilia morphology despite Odad1 KD. (I-K) GFP-Dnai2 is restored to Odad1 deficient motile cilia after dose gradient ectopic expression of Myc-Odad1. (C′-D’) Merged channels show membrane (magenta) and Dnai2 (green) labeling in rescued MCCs. (E) Quantification of mean±s.d. of normalized GFP-Dnai2 fluorescence (see Methods) in axonemes of control (n=141), Odad1 KD (n=137), 350 pg rescue (n=107) and 500 pg rescue (n=105) MCCs. *N*>25 cells from nine embryos across three experiments for all conditions. (F,G) GFP-Dnal4 labeling of motile axonemes in control (F) and Odad1 KD (G) MCCs. GFP-Dnai2 shows normal localization in control MCCs but is lost from axonemes in Odad1 KD MCCs. (F′,G′) Merged channels show membrane (magenta) and Dnal4 (green) labeling for cells in panels F and G, respectively. (H) Quantification of normalized GFP-Dnal4 mean intensity along axonemes in control (n=53), Odad1 KD (n=54) and rescue (n=55) MCCs. N>25 cells from nine embryos across three experiments for all conditions. Loss of GFP-Dnal4 is restored after ectopic expression of Odad1. (I) GFP-Wdr78 labeling of IDA-f subunits shows normal localization in motile axonemes. (J) GFP-Wdr78 remains localized to motile cilia in Odad1 KD MCCs. Merged channels (bottom row) show normal cilia morphology in both control and Odad1 KD MCCs, with membrane (magenta) and Wdr78 (green) labeling. (K) GFP-Ttc29, an IDA-d subunit, also labels ciliary axonemes in control MCCs. (L) Odad1 KD does not change the localization pattern of Ttc29. (M,N) Graphs showing mean intensity of normalized IDA subunits Wdr78 (M) and Ttc29 (N) in control and Odad1 KD MCCs. Wdr78 control axonemes quantified n=102 and Odad1 KD n=121. Ttc27 control n=108 and Odad1 KD n=141. Data are presented as mean ± SD with error bars representing independent biological replicates. Statistical significance was determined using a non-parametric Mann–Whitney *U*-test for two-group comparisons and a one-way ANOVA for comparisons involving more than two groups. ***p<0.001; ns, p>0.05. Scale bars: 10 µm.

Inner dynein arms (IDAs) are a class of axonemal dyneins that control ciliary waveform but are not directly associated with ODA-DC [17], [44]. Importantly, IDA were reported unaffected in ODAD1 patients [18], [19], [23]. To assess specificity of Odad1 in the deployment of ODAs to microtubules, we investigated the effect of Odad1 loss on IDA deployment to MCCS. We found that Odad1 knockdown did not disrupt the axonemal localization of the IDA-f subunit GFP-Wdr78 (Fig. 2I-J, I’J’, M) nor the IDA-d subunit GFP-Ttc29 (Fig. 2K-L, K’L’, N). These findings indicate that Odad1 is specifically required for deploying ODA, but not IDA, subunits to the axoneme in *Xenopus* MCCs.

### Odad1 is required for deployment of ODA-DC subunits to the axoneme

The outer dynein arm docking complex (ODA-DC), composed of Odad1-Odad5, is required to anchor outer dynein arms (ODAs) along the axoneme, and loss of individual ODA-DC subunits has been linked to PCD. To test whether Odad1 is required for localization of the other ODA-DC subunits to axonemes, we expressed GFP-tagged Odad2, Odad3, Odad4, or Odad5 in *Xenopus* MCCs and compared their ciliary localization in control vs Odad1 KD MCCs by confocal microscopy.

In control MCCs, all four GFP-tagged subunits localized robustly along the length of the ciliary axoneme (Fig. 3A-D, A′-D′). Following Odad1 KD, axonemal localization of each subunit was markedly reduced (Fig. 3 E-H, E′-H′), indicating that Odad1 is required for their deployment to cilia. Quantification of normalized GFP mean intensity confirmed a significant decrease for all four subunits in Odad1 KD MCCs relative to controls: Odad2 (Fig. 3I), Odad3 (Fig. 3J), Odad4 (Fig. 3K), and Odad5 (Fig. 3L).

**Figure 3.**
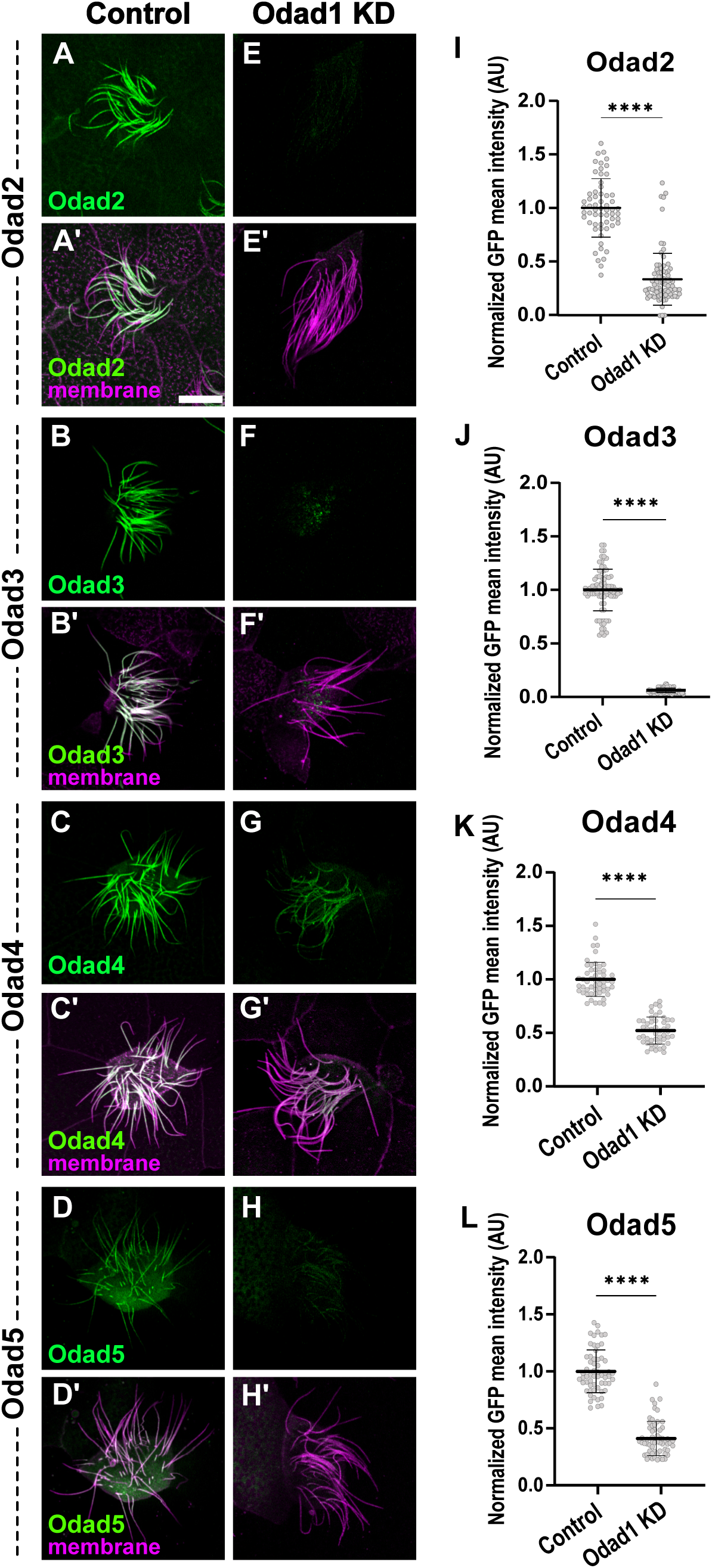
Odad1 is required for deployment of ODA-DC subunits in *Xenopus* MCC. (A–D) Representative confocal images of GFP-Odad2 (A), GFP-Odad3 (B), GFP-Odad4 (C), and GFP-Odad5 (D) (green) decorating the ciliary axoneme in control multiciliated cells (MCCs). (A′–D′) Merged images of the corresponding panels above, showing membrane (magenta) and ODA-DC subunit (green) localization. (E–H) GFP-Odad2 (E), GFP-Odad3 (F), GFP-Odad4 (G), and GFP-Odad5 (H) localization to MCCs is reduced following Odad1 knockdown (KD), compared to controls. (E′–H′) Merged images of the corresponding panels above. (I–L) Quantification of normalized GFP mean intensity for Odad2 (I), Odad3 (J), Odad4 (K), and Odad5 (L) in control versus Odad1 KD MCCs. Each point represents one cilium (Odad2: control n=60 and Odad1 KD n=82, Odad3: control n=90 and Odad1 KD n=94, Odad4: control n=53 and Odad1 KD n=48 and Odad5: control n=60 and Odad1 KD n=60). N>25 cells from nine embryos across three independent experiments were analyzed per condition; bars show mean ± SD. Statistical significance was determined by one-way ANOVA for multiple comparisons. ****P<0.0001 Scale bars: 10 µm.

The magnitude of reduction varied across subunits, with Odad3 showing the most severe loss of ciliary signal upon Odad1 KD (Fig. 3 F-F’, J). Notably, the degree of decrease correlated with each subunit’s proximity to Odad1 in the previously reported cryo-EM structure [13], consistent with Odad1 and Odad3 being closely positioned within the complex. Furthermore, cryo-EM models form axonemes from organoids derived from ODAD1 patients have also shown that loss of ODAD1 is enough for the loss of the entire ODA-DC density [17]. Together, these data suggest that Odad1 acts upstream of the other ODA-DC components and is essential for their trafficking or stable docking to the axoneme, positioning Odad1 as a core organizer of ODA-DC assembly in vertebrate MCCs.

### Disease-associated *ODAD1* variants are destabilizing, loss-of-function alleles

To determine whether patient-derived ODAD1 alleles are functionally deleterious, we generated *Xenopus laevis* Odad1 constructs harboring truncating mutations at residues homologous to two ciliopathy-associated frameshift alleles, p.L105fs and p.P236fs, yielding Xla p.T140* and Xla p.I270*, respectively (Fig. 4A). Both alleles fall within or immediately downstream of predicted coiled-coil domains, which based on mammalian cryo-EM structure are important for its interaction with microtubules, hence, proper ODA-docking complex attachment.

**Figure 4.**
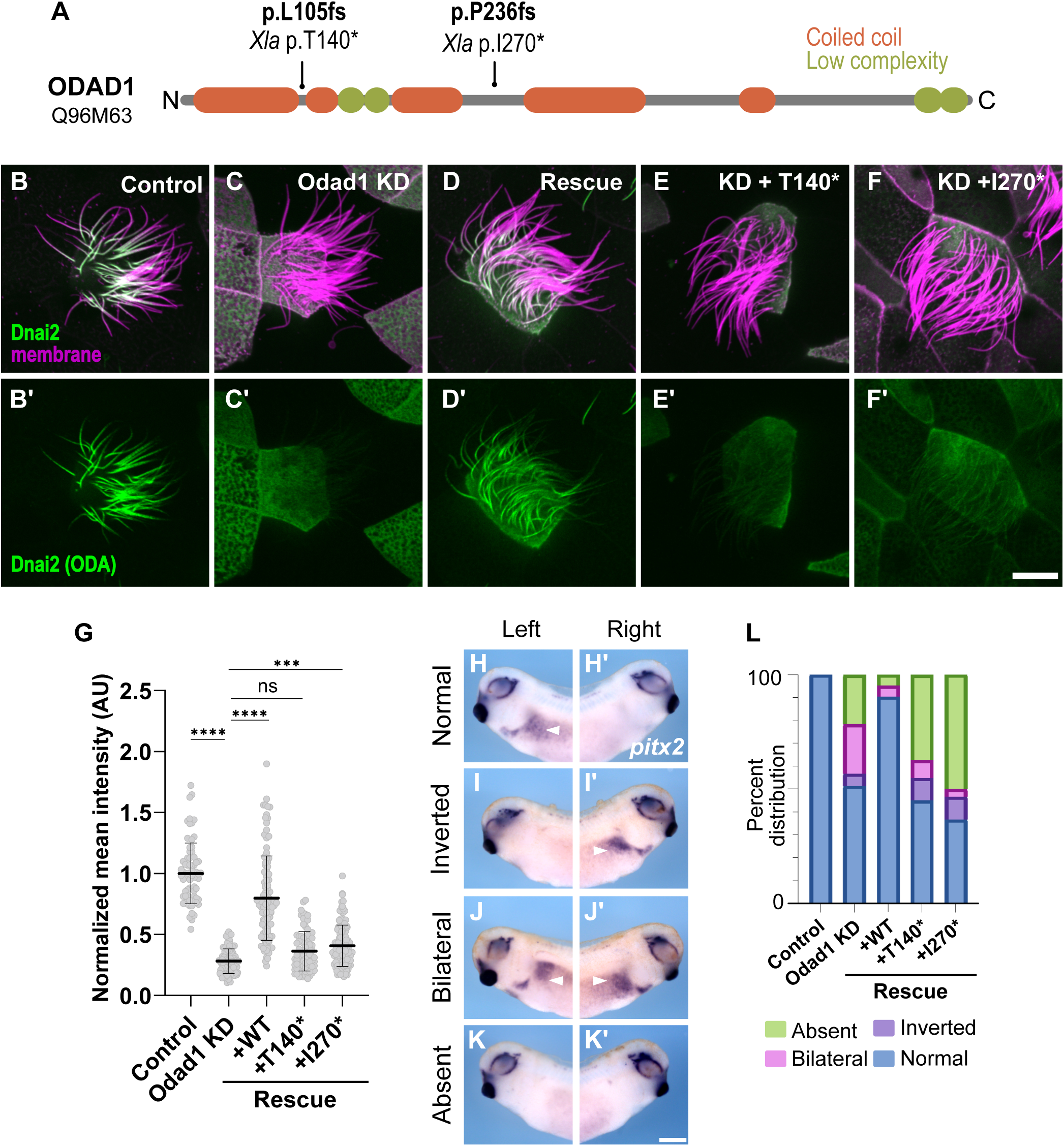
Patient-derived ciliopathy alleles fail to restore ODA and left-right asymmetry defects. (A) Schematic of ODAD1 (Q96M63) showing coiled-coil (orange) and low-complexity (green) domains. Positions of patient frameshift alleles (p.L105fs, p.P236fs) and their corresponding truncating mutations generated at homologous residues in *Xenopus laevis* Odad1 (Xla p.T140*, Xla p.I270*) are indicated. (B-F) Multiciliated cells from *Xenopus* embryos showing localization for the ODA subunit Dnai2 (green) and membrane (magenta): control (B), Odad1 knockdown (KD) (C), KD rescued with wild-type Odad1 (D), KD + Odad1 p.T140* (E), and KD + Odad1 p.I270* (F). (B′-F′) Dnai2 channel alone for each condition above. Odad1 KD reduces ciliary Dnai2 signal; wild-type Odad1 restores it, whereas the two patient-equivalent truncating alleles fail to rescue. Scale bar: 10 µm. (L) Quantification of normalized mean Dnai2 intensity per cell across conditions in (B-F). Each point represents an individual cilium, in control (n=73), Odad1 KD (n=93) and rescue +WT (n=109), +T140* (n=85), +I270* (n=125) MCCs. N>25 cells from nine embryos across three experiments for all conditions; bars show mean ± SD. \*\**P* < 0.001, \*\*\*\**P* < 0.0001, ns = not significant. (H-K′) Whole-mount in situ hybridization for *pitx2* in stage 28 embryos, shown from left (H-K) and right (H′-K′) views, illustrating the four scored left-right patterning outcomes: normal (left-restricted expression, H/H′), inverted (right-restricted, I/I′), bilateral (J/J′), and absent (K/K′). Scale bar: 25 mm. (L) Stacked bar graph showing the percent distribution of *pitx2* expression patterns (normal, bilateral, inverted, absent) in control, Odad1 KD, and KD embryos rescued with WT, p.T140*, or p.I270* Odad1. Patient-equivalent alleles fail to restore normal left-right patterning to control levels, phenocopying Odad1 KD.

We first asked whether these patient-equivalent alleles could restore ODA deployment in Odad1-depleted MCCs. Expression of GFP-tagged Xla p.T140* and Xla p.I270* constructs resulted in failure to localize to axonemes in MCCs, in contrast to wild-type Odad1, which localized properly to axonemes (Fig. S3). As previously observed, Odad1 KD markedly reduced ciliary localization of the ODA subunit Dnai2 relative to control MCCs, and this defect was rescued by re-expression of wild-type Myc-tagged Odad1 (Fig. 4 B-D, B’-D’, G). In contrast, expression of either Myc-tagged Odad1 p.T140* or p.I270* in the KD background failed to restore Dnai2 signal to cilia, with intensity levels indistinguishable from Odad1 KD alone (Fig. 4E-F, E’-F’, G). These results indicate that both patient-derived truncations make Odad1 non-functional for ODA docking.

We next asked whether these alleles could rescue the left-right patterning defects caused by Odad1 loss, scored by *pitx2* expression as previously described (Fig. 1A-D, A’-D’). Co-injection of wild-type Odad1 largely restored normal, left-restricted *pitx2* expression, whereas embryos co-injected with Odad1 p.T140* or p.I270* remained predominantly abnormal, with *pitx2* phenotype distributions closely resembling Odad1 KD alone (Fig. 4L). Importantly, western blot analysis of Myc-tagged constructs showed that, even at rescue doses (500 pg), both truncated proteins were unstable and expressed at lower levels compared to wild-type Odad1, explaining their failure to rescue the knockdown phenotype (Fig. S3). Together with the ciliary Dnai2 rescue data, these findings demonstrate that both patient-derived ODAD1 truncating alleles are loss-of-function variants that fail to support ODA assembly and downstream left-right patterning, consistent with their pathogenicity in ciliopathies.

## Discussion

ODAD1 is a core component of the pentameric ODA-DC [13], and variants in this gene are known to cause PCD frequently presenting with laterality defects [18], [19], [20], [21], [22], [23], [24]. Here, we investigated the function of Odad1 *in vivo* by combining knockdown and rescue approaches with analysis of patient-derived disease alleles across two ciliated cell types in *Xenopus* embryos: nodal cilia at the embryonic GRP and MCCs of the embryo epidermis. Our results show that Odad1 is required for LR asymmetry establishment, as reflected by disrupted *pitx2* expression (Fig. 1D-H) and gut looping (Fig. 1A-C), and for deployment of ODAs and ODA-DC subunits to MCC axonemes (Fig. 2, 3), which results in downstream fluid flow anomalies (Fig. S2). We further find that two previously uncharacterized patient-derived truncating alleles are unstable, loss-of-function variants that fail to support ODA docking or LR patterning (Fig. 4).

Odad1 has not previously been reported in the axonemes of 9+0 motile cilia, despite its established role in human laterality. In the frog GRP, GFP-Odad1 localized along the full length of nodal cilia axonemes and was additionally enriched at the ciliary base, extending around the basal body (Fig. 1I-L). Odad1 localization to the 9+0 nodal cilia axonemes is consistent with our findings in 9+2 MCCs cilia (Fig. 2A-H, S3A-A’) and with structural data from 9+2 cilia in the bovine airway [13]. However, it contrasts with a report of Odad1 restricted to basal bodies and absent from axonemes in zebrafish Kupffer’s vesicle and spinal canal [45]. Several reports suggest that the axonemal localization we observe is correct. Other ODA-DC components localize to 9+0 axonemes, for instance Odad2 in the zebrafish Kupffer’s vesicle and spinal canal [44], Odad4 and Odad5 in the mouse node cilia [28], [46]. The zebrafish Odad1 result may reflect species or cilia-type differences, or the detection method, since immunolabeling of fixed, epitope-tagged proteins and live imaging often give different localization patterns for ciliary proteins. Axonemal Odad1 localization is also consistent with laterality phenotypes we and others report [14], [15], [27], [28] and with ODA-driven beating in 9+0 cilia, which is disrupted in ODA-DC mutants [28], [47]. Moreover, Odad1 loss reduced axonemal Odad3 in nodal cilia (Fig. 1M–O). We therefore conclude that Odad1 acts in the axoneme, where it supports docking of downstream ODA-DC components, and provides a cellular basis for the LR defects seen upon its loss.

Our emerging understanding of the mechanisms of cytosolic transport of ODAs has opened an avenue of new questions including, including how the ODA-DC is assembled and trafficked in the cytosol. Pre-assembly of ODA-DC in motile cilia remains largely unexplored. Data from unicellular ciliated organisms, have shown that ODAs and ODA-DC are deployed to ciliary axonemes independently [26], [48]. Our MCC data shows that loss of Odad1 dramatically decreases localization of other ODA-DC subunits, specifically Odad3. These results could suggest a role in pre-assembly of ODA-DC. Furthermore, given that both Odad1 and Odad3 are enriched at the ciliary base of nodal cilia in the *Xenopus* GRP, we hypothesize that the ODA-DC may have additional roles in these cells; this accumulation could suggest a recruitment or pre-assembly site for the ODA-DC. Further experiments to test this hypothesis are warranted.

Despite the identification of numerous ODAD1 ciliopathy-associated variants, only six variants have been experimentally validated *in vitro* [11–15] leaving most unresolved. The two patient-equivalent truncating alleles we modeled, Odad1 p.T140* and p.I270*, both fall within or immediately downstream of predicted coiled-coil domains implicated in microtubule attachment, were unable to localize to axonemes. Expression levels of both variants were markedly reduced compared to wild-type protein, and the mutant proteins consequently failed to rescue either ODA docking or LR patterning defects (Fig. 4, Fig. S3). These results indicate that these alleles act through protein destabilization rather than a subtler disruption of a specific interaction interface, supporting a straightforward loss-of-function mechanism underlying ODAD1-associated PCD. Systematic characterization of ODAD1 variants for localization and function, prioritized based on structural context and evolutionary conservation — particularly at interfaces with microtubules and ODA-DC subunits — is crucial to further understand mechanisms underlying ODAD1-associated ciliopathies.

In summary, our results support a model where Odad1 functions as a central organizer of ODA-DC assembly required for both 9+0 cilia and LRO-driven laterality and 9+2 cilia and MCC ciliary motility in vertebrates and demonstrates that patient-derived truncating alleles are destabilizing loss-of-function variants. By linking ODA-DC assembly mechanisms to LR patterning defects and to human disease alleles *in vivo*, this study provides a framework for further dissecting the hierarchical assembly of the ODA-DC and for testing additional ODAD1 patient variants in a vertebrate context.

**Supplemental Figure 1. Odad1 knockdown RT-PCR**

Validation of Odad1 splice-blocking morpholino efficiency by RT-PCR. Gel image of RT-PCR of *odad1* and *odc1* mRNA levels in wildtype control (WT), Odad1 MO 10 ng and 20 ng injected embryos.

**Supplemental Figure 2. Flow trace of fluorescent bead flow stage NF 28 Xenopus laevis embryos.**

Bead trajectories over 60 frames per second using a temporal-color code, mapping frame number to a fire color scale from frame 0 (Blue) to frame 60 (white). Embryo stage NF 28 showing control (A) and Odad1 KD (B) bead trajectories.

**Supplemental Figure 3. Odad1 patient alleles localization to MCCs and protein abundance.**

*Xenopus* variants (human): T140* (L105*) and I270* (P236*). (A) Wildtype localization of Odad1 in MCCs. Odad1 (green) localizes along the axoneme except the most distal end of the cilium. (B) Expression of the T140* and (C) I270* allele in MCCs disrupted its localization to axonemes compared to that in controls. (A’-C’) Merged image of Odad1 (green) and a membrane marker (magenta) in control (A′), T140* (B’) and I270* (C′) MCCs. (B) Western blot showing protein levels for indicated disease alleles when expressed in *Xenopus* embryos. Western blot of total protein from N=30 embryos NF 25, injected with 500pg of GFP-Lrrc56 WT, T140*, and I270*. Anti Myc 1:1000 and Anti B-actin housekeeping control (1:10,000).

## Acknowledgements

We thank Xenbase and the European Xenopus Resource Centre for providing *Xenopus* resources.

