## Supplementary figures and images for "Odad1 is Required for Deployment of Axonemal Outer Arm Dyneins and Embryonic Left-Right Patterning"

### Supplemental Figure 1

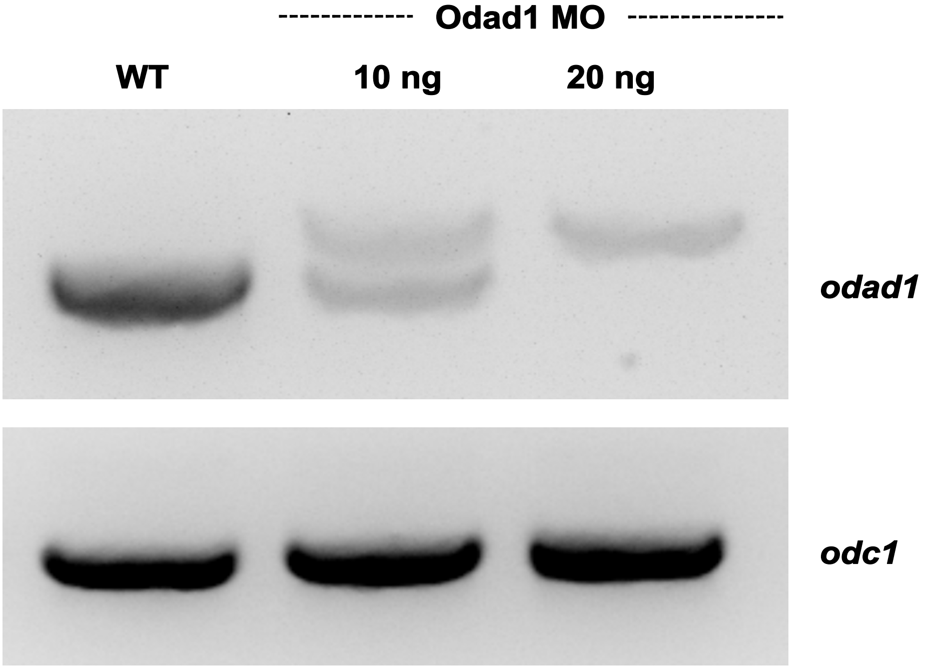

### Supplemental Figure 2

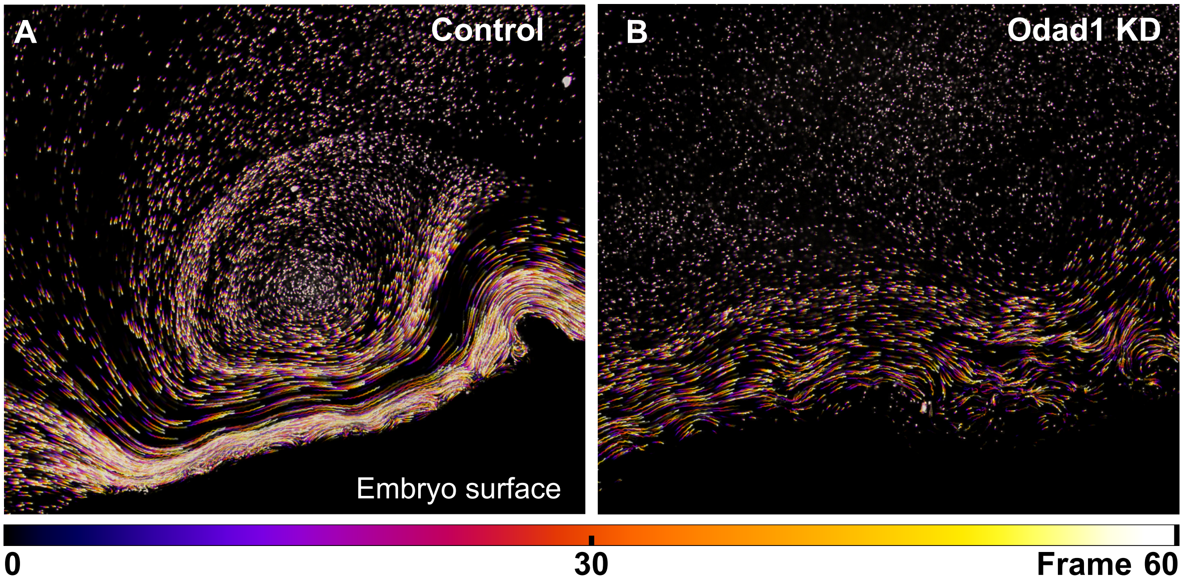

### Supplemental Figure 3

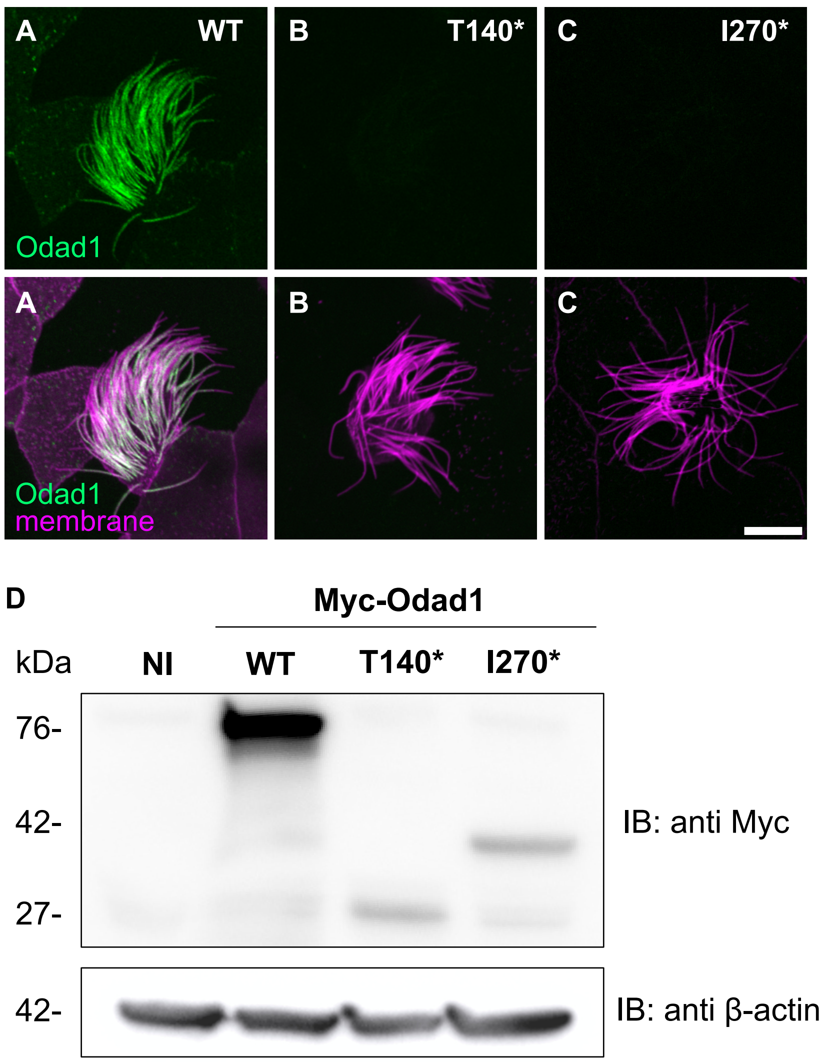
